# Predicting Protein Thermostability Upon Mutation Using Molecular Dynamics Timeseries Data

**DOI:** 10.1101/078246

**Authors:** Noah Fleming, Benjamin Kinsella, Christopher Ing

## Abstract

A large number of human diseases result from disruptions to protein structure and function caused by missense mutations. Computational methods are frequently employed to assist in the prediction of protein stability upon mutation. These methods utilize a combination of protein sequence data, protein structure data, empirical energy functions, and physicochemical properties of amino acids. In this work, we present the first use of dynamic protein structural features in order to improve stability predictions upon mutation. This is achieved through the use of a set of timeseries extracted from microsecond timescale atomistic molecular dynamics simulations of proteins. Standard machine learning algorithms using mean, variance, and histograms of these timeseries were found to be 60-70% accurate in stability classification based on experimental ΔΔ*G* or protein-chaperone interaction measurements. A recurrent neural network with full treatment of timeseries data was found to be 80% accurate according the F1 score. The performance of our models was found to be equal or better than two recently developed machine learning methods for binary classification as well as two industry-standard stability prediction algorithms. In addition to classification, understanding the molecular basis of protein stability disruption due to disease-causing mutations is a significant challenge that impedes the development of drugs and therapies that may be used treat genetic diseases. The use of dynamic structural features allows for novel insight into the molecular basis of protein disruption by mutation in a diverse set of soluble proteins. To assist in the interpretation of machine learning results, we present a technique for determining the importance of features to a recurrent neural network using Garson’s method. We propose a novel extension of neural interpretation diagrams by implementing Garson’s method to scale each node in the neural interpretation diagram according to its relative importance to the network.

## I. Introduction

Advances in genetic sequencing technologies and algorithms are enabling the use of genetic information as a diagnostic tool for clinicians treat patients. However, it still remains a significant challenge to identify disease-causing mutations among the much more commonly occurring neutral mutations found in human populations with high accuracy. This is largely due to the existence of an estimated 10,000 nonsynonymous variations in each human genome, which has prevented experimental characterization using existing methods [1]. It is for this reason that a wide-range of computational tools have emerged to assist in the annotation of mutations. In many cases, such tools have a related application for the optimization of protein thermostability in the field of protein engineering.

In this report, we begin by discussing several ways of characterizing the effect of mutations in protein-coding genes, as well as existing experimental and computational approaches designed to predict these effects. To this end, we discuss the value of protein crystal structure and homology models along with how they have been utilized in the literature. Next we describe the importance dynamic structural features and how they may provide new insight into the effect of mutation on structure.

### A. Clinical Significance of Understanding Genetic Alterations

Humans are roughly 99.5% identical in genetic make up, but this critical 0.5% largely determines how humans develop diseases and respond to pathogens, chemicals, drugs, vaccines, and other agents [2]. This genetic difference between individuals results from alterations including single nucleotide variants (SNVs), small insertions or deletions (indels), gene fusions, copy number variations, and large chromosomal rearrangements. In this report, we examine the effect of SNVs in protein coding results that result in changes to an amino acid in a resultant protein, referred to as a missense mutation. In a number of well-studied diseases, missense mutations result in a loss of function of protein coding genes. Structural modifications through mutation may impair normal protein function or prevent protein folding entirely. Numerous mendelian diseases, such as cystic fibrosis [3] and muscular dystrophy [4], have been linked to deleterious mutations in human proteins (in the cystic fibrosis transmembrane receptor and dystrophin proteins, respectively). This relationship was established based on targeted biochemical, structural, and genetic experiments. However, given that over 64 million genetic mutations have been identified in humans [5], the experimental study of each mutation is not possible due to high costs and technical difficulty. To overcome these barriers, scientists have developed proteome-wide computational and statistical tools for the identification of mutations that negatively affect protein function. A high-accuracy approach of this nature may greatly impact the diagnosis, prevention, and treatment of diseases resulting from missense mutations.

### B. Measuring the Effect of Missense Mutations

There are numerous ways to measure the effect of missense mutations. Most commonly, the effect of mutation is characterized by its deleteriousness or pathogenicity, where the former refers to the disruption of protein structure (resulting in lowered stability or misfolding) and the latter refers to an experimental correlation with disease. One way to quantify mutation deleteriousness is to measure its ΔΔ*G* of folding. This refers to the change in energy involved with folding the protein from an extended state to its native state, with or without a single-point mutation, and is a common metric for stability. One of the largest databases of experimentally measured ΔΔ*G* values is known as ProTherm, and is used extensively in this work [6]. Similarly, computational techniques such as alchemical free energy calculations may be used to compute a related ΔΔ*G* quantity to allow for the inference of protein stability [7], [8]. ΔΔ*G* of folding is expected to change based solely on the physicochemical properties of the exchanged amino acids (charge, size, and other chemical properties of their respective side chains), but even more complicated effects could occur. As the process of protein folding may involve non-native interactions not readily apparent in existing protein models, it is difficult to predict how a mutation may influence stability. In an extreme case, a mutation may result in large-scale conformational change that may disrupt the fold of the protein, potentially resulting in disease due to the aggregation of misfolded protein (thought to occur in prion disease). Interestingly, even subtle conformational changes that may not significantly alter stability could still result a loss-of-function of the protein and disease. A study of protein-chaperone interactions involving proteins with disease causing mutations found that the majority of mutations did not impair protein folding or stability [9]. Chaperones are proteins that assist in the repair of misfolded proteins and specific biochemical assays can be used to measure protein-chaperone interactions which could also be used to measure protein stability upon mutation [10]. This underscores the importance of selecting a metric for the effect of mutation, not only for the interpretation of results with respect to disease, but for comparisons to other methods in the literature. In this work, we performed classification in order to predict neutral or deleterious changes to protein stability using either the sign of a ΔΔ*G* value or the presence of protein-chaperone interactions. We expect that a significant proportion of ΔΔ*G* values close to zero may be mislabelled in our dataset and that the sensitivity of protein-chaperone interaction assays may present a source of error in our study.

### C. Structural Characterization of the Human Genome

The Protein Data Bank contains over 100,000 protein structures across a wide range of organisms. From surveys of all proteins in this ensemble, researchers have identified that much of the proteome consists of evolutionarily conserved domains with relatively few unique three-dimensional folds. A recent analysis by Perdigao et al. suggests that for a major annotated database of 546,000 protein sequences (Swiss-Prot), 56% of the proteome in eukaryotes could be matched to a homologous protein with known structure [11]. The determination of structure based on homologous proteins is facilitated by homology modelling using software like Rosetta or MODELLER after performing sequence alignment [12], [13]. Although a large percentage of eukaryotic proteins are still unknown, and may correspond to completely unknown folds, advances in structure determination methods are rapidly accelerating the discovery of protein structures. This suggests that proteome-wide structural analysis of proteins may become increasingly useful for the characterization of protein stability and genetic diseases, especially when studying a specific subset of proteins with diverse folds.

### D. Approaches to Predict Protein Stability

A number of approaches have been utilized to predict protein stability upon mutation. Some of most widely utilized computational techniques employ protein sequence data [14]. Both conserved and non-conserved regions exist in protein sequences across multiple organisms. By examining protein sequence throughout evolution, it can be inferred that mutations in conserved regions may result in a loss of stability or function of the protein, whereas mutations in non-conserved regions may have little effect. However, conservation driven models like this have been known to yield true positive rates less than 50% in less conserved regions, encounter difficulty in diagnosing benign variations in conserved positions, and have poor accuracy for single nucleotide variants associated with complex diseases [15]. With the increasing availability of structural data for protein-coding genes, new approaches are combining sequence and structure based data in order to improve protein stability predictions.

All of the following approaches employ machine learning to predict protein stability upon mutation. These approaches include neural networks [16], [17], random forests [18]–[20], decision trees, [21], [22] and support vector machines [23]–[26]. In a study by Jia et al. [27], five supervised machine learning methods (support vector machines, random forests, neural networks, the naive Bayes classifier, and K-nearest neighbours) along with partial least squares regression were benchmarked for performance in predictive modelling of protein stability. Some of these studies report a high degree of success in predicting mutational effects with either binary or ternary classification (binary classification as stabilizing or destabilizing, or ternary classification as stabilizing, no effect, or destabilizing). Several studies train regression-based models in order to obtain quantitative values for ΔΔ*G* to be compared to ΔΔ*G*_*exp*_ using Pearson correlation. A comparison of these studies is often made difficult by differences in training/validation/testing datasets, training methodology, dataset sizes, and hyperparameter optimization. Nonetheless, Jia et al. [27], perform many methodological variations and report the highest binary classification accuracy (0.90) with Rosetta energy features [12]. Similarly, Berliner et al. [21] report the highest regression correlation between predicted and experimental values (0.77) combined with FoldX energy features [28]. Both studies utilize data from the ProTherm database as targets for machine learning.

Several studies have employed homology modelling to construct a large dataset of protein structures which could be used to derive structural features [21], [23], [27]. Previous attempts at using protein structure were limited by the low number of human proteins available in the Protein Data Bank. Upon constructing a large ensemble of human protein homology models, structural features were extracted, which include secondary structure type, solvent accessible surface area, charge environment, and other metrics. These features were then combined with energy-function derived features using the FoldX or Rosetta algorithms [12], [28], in addition to sequence-based features. These approaches were found to be effective at protein stability prediction, but still suffer from some limitations. In Berliner et al. [21], several of the most predictive features in this approach were derived from the FoldX algorithm, of which these features were likely constructed using empirical data from the ProTherm database. As such, it is expected that the agreement with experimental values in that study may be overestimated due to overfitting. All of these studies utilizing protein structure rely heavily on homology models of wildtype and mutant proteins, using little to no structural validation [21], [23], [27]. The assumption of these approaches is that single-point mutations do not largely alter the folding of the protein, even though this has been observed within a subset of disease-causing mutations in humans [9]. Lastly, while Baugh et al. generated as many as 50 structures for wildtype and mutant proteins using Rosetta [23], there is no explicitly dynamic information in these models. It is possible that crystal structures and homology models created may not correspond to the most probable physiological state of the protein. These issues have motivated us to include dynamic structural data from protein simulations in order to improve protein stability predictions.

### E. Protein Dynamics

Proteins are known to adopt multiple distinct conformational states to perform specific biological functions. These conformational changes facilitate interactions with water, ions, other proteins, and other biomolecules, all of which can not be directly inferred from protein sequence alone. Although static structural snapshots of proteins provide the basis for establishing a structure and function relationship, additional site-specific experimental studies are required to verify our understanding of protein dynamics. One of such techniques involves constructing a computational model of a protein and utilizing physics-based algorithms to sample conformations available to the protein (Figure 1). State-of-the-art protein simulations have some known limitations arising from a combination of systematic (accuracy in force fields) and statistical errors (insufficient statistical sampling), but are widely considered accurate enough to reproduce many experimental measurements.

**Fig. 1.**
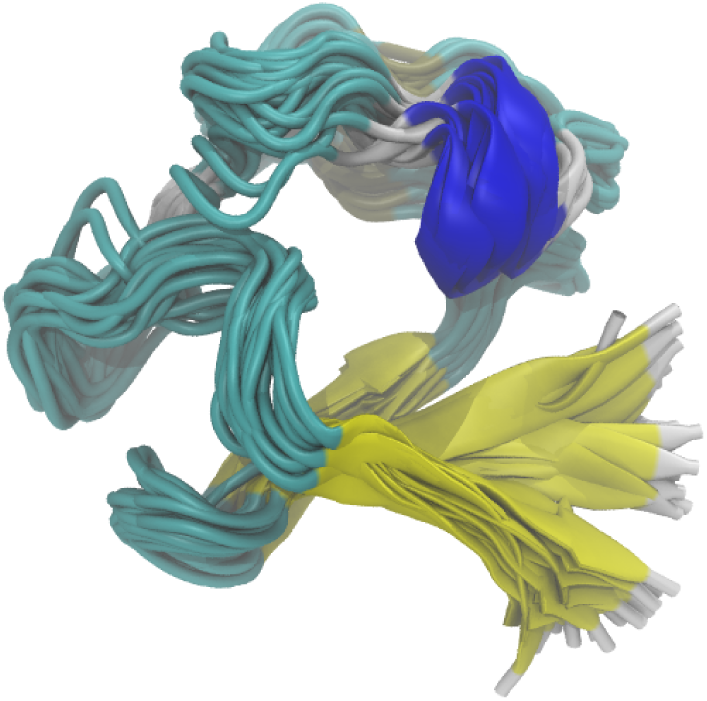
Rendering of multiple time frames in simulations of the protein rubredoxin (PDB: 1BFY). Protein is colored based on secondary structure.

The effect of protein mutations can be studied using molecular simulations in a number of ways. In a small number of cases, protein structures have been solved with and without mutations, and thus molecular models could be constructed and simulations performed on both proteins, respectively. In the remaining cases where only wildtype protein structures are available, homology models can be constructed that introduce a single-point mutation. However, the latter approach may require extensive validation as some mutations have been known to cause unfolding, side-chain repacking, or even the emergence of new structures. In this case, molecular simulations are frequently employed in order to sample a protein conformation more representative of the physiological state, under the assumption that the mutation does not severely effect protein folding. In order to reduce the dependence of our approach on the existence of high quality homology models of mutants, we propose that machine learning may be used to study the effect of mutations without explicitly modelling and simulating them. In this study, we assume the effect of a missense mutation may be broadly inferred from the dynamic fluctuations of the protein in its wildtype form. The use of protein dynamics to study mutational effects has been performed on a small scale, but without machine learning approaches [29], [30]. By examining the conformational fluctuations of a protein over time, we propose that machine learning approaches may be utilized to predict the effect of mutation. This may be achieved by extracting information on both the dynamic environment of a prospective mutation, physicochemical properties of the newly introduced amino acid. For example, a region within the overall topology of a protein that may be considered a flexible hinge or linker to facilitate conformational change, would be disrupted by certain mutations depending on its hydrophobicity, size, or charge, ultimately causing loss of protein stability. The efficacy of this approach is dependant on the correct featurization of the protein and site of mutation.

### F. Machine Learning using Timeseries Data

The primary goal of our approach is to better predict how a specific mutation impacts a protein’s stability, which is represented by atomic trajectories over time. In this report we investigate three approaches for learning to classify timeseries data. As a naive approach, the mean and variance is extracted for each timeseries feature, and used as features to multiple machine learning methods. Following this, we utilized a modified “bag-of-words” model, where the frequency of discretized timeseries values are used as features to train machine learning classifiers. For example, we may featurize the site of mutation by the number of water molecules found in it’s vicinity. Using a modified “bag-of-words” model, the bins represent a discrete number of water molecules near the mutation site, and the bars represent the frequency of this solvation state. For these features lacking explicit timeseries information, we aim to explore several machine learning approaches implemented in the scikit-learn package [31]. We did not calculate the classification performance of simple machine learning methods by sampling multiple static snapshots along our trajectory.

We consider recurrent neural networks (RNNs) as the primary model of interest to our study for several reasons. Unlike many of the other methods for handling temporal data, recurrent neural networks are not restricted to a fixed size dependency, and in theory are able to cover arbitrary length dependencies throughout the input sequence [32], [33] which are likely to occur during protein simulations.

Recurrent neural networks have been employed to achieve state of the art performance on a variety of problems throughout many domains. These include problems in text classification [34], speech recognition [35], modelling genetic regulations inside cells [36], language modelling [37], and machine translation [38]. Although neural networks are typically employed only for data-abundant tasks, several results have shown that carefully regularized neural networks can perform quite well on small datasets [39] [40]. Menkovski et al. [40] study whether deep neural networks can be trained to perform well in data-limited scenarios, focusing on the task of identifying anatomy within x-ray images. While we intend to use traditional machine learning approaches as well, to our knowledge the use of RNNs is a novel approach to predicting the stability of mutations in protein data.

## II. Recurrent Neural Networks

Recurrent neural networks (RNNs) were introduced as a means to overcome the inability of feed-forward neural networks to handle temporal data, in which inputs may be sequences of variable length and points within the temporal sequence may depend on each other. RNNs extend regular neural networks by allowing them to extract temporal dependencies between examples within a sequence. Given a timeseries *x* = *x*_1_*,…,x*_*T*_, each element *x*_*t*_ is fed sequentially into the neural network. Intuitively, each hidden unit within an RNN has a *memory* which allows it to remember important features of the portion of the timeseries which it has seen, and discover temporal correlations between events in the data.

More specifically, each hidden unit of a Recurrent Neural Network is a *recurrent unit*, it contains a recurrent state whose activation depends on the input to that hidden unit, as well as the activation of the recurrent state from the previous step; an illustration of this is given in figure II [41]. Precisely the recurrent state of the hidden unit *h* at time *t* is given by

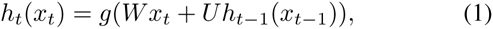

where *W* and *U* are weights on the edges, and *g* is some smooth bounded function. Several choices arise for the output of an RNN.

**Fig. 2.**
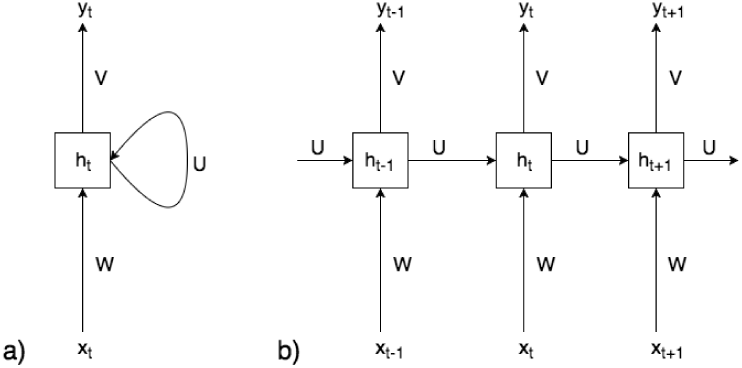
(a) The structure of a recurrent unit in an RNN. (b) The recurrent unit unrolled over time.

The recurrent unit can either produce an output *y*_1_ = *h*_1_(*x*_1_),…,*y*_*t*_ = *h*_*t*_(*x*_*t*_) for each entry *x*_1_,…,*x*_*T*_ in the time series as is done in the *many-to-many model*, produce only a single output *y*_*T*_ = *h*_*t*_(*x*_*T*_) after every entry of the time series has been seen as is done in the *many-to-one* model, or some intermediate between the two. The recurrent neural networks which we designed are of the first two types.

Although the distance of temporal dependencies captured by RNNs do not have an explicit limitation, equation 1 shows that the dependency on an example *x*_*i*_ decreases exponentially as we move away from *i* in the sequence. Therefore, in reality RNNs only have *short-term memory*.

### A. Long Short-Term Memory

To address the issue of handling long-term dependencies, Hochreiter et al. [42] introduce a more involved recurrent unit known as the Long Short-Term Memory (LSTM) unit. The idea behind the LSTM is a recurrent unit which is able to decide what to remember and what to forget, allowing it to handle long-term dependencies that the regular recurrent units could not. The LSTM is composed of a *memory cell state*, *c*_*t*_ which contains the information remembered by the LSTM unit at time *t* in the form of a self-recurrent connection. Information is added and removed from the memory cell state by a series of gates.

Let *W*_*f*_, *W*_*i*_, *W*_*c*_, *W*_*o*_, *U*_*f*_, *U*_*i*_, *U*_*c*_, *U*_*o*_ be weight matrices and *b*_*f*_, *b*_*i*_, *b*_*c*_, *b*_*o*_ be bias vectors. Given the the next input *x*_*t*_ in a timeseries *x*_1_,…,*x*_*T*_ and the output of the LSTM unit at the previous input in the sequence, *h*_*t*−1_, the forget gate *f*_*t*_ determines what information should be remove from the memory cell state,

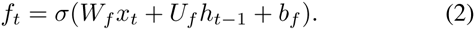

The input gate *i*_*t*_ then decides which values should be updated in the memory cell state,

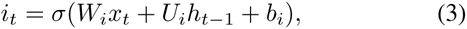

and a candidate update 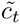 is created

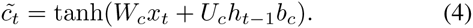

The partially forgotten previous memory cell state *f*_*t*_*c*_*t*_−_1_ is combined with the to-be-updated values of the candidate state 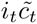 to form the new memory cell state

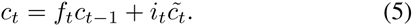

Finally the values of the memory cell state which the LSTM unit will output is decided by the output gate based on the previous output of the LSTM *h*_*t*_−_1_ and the input

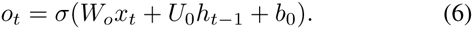

The output of the LSTM unit is calculated as

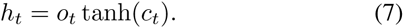

### B. Bidirectional Recurrent Networks

Siwei et al. [34] gave state of the art performance for problems of text classification by creating a neural network with the intuition of representing each word in a text with its context within that text. To do this they use a bidirectional recurrent neural network to capture information about the words which appear before and after it. Introduced by Schuster et al. [41], a bidirectional recurrent neural network is a generalization of a recurrent network in which each recurrent node has both a forwards-in-time and backwards-in-time recurrent loop. This can be seen in figure 3. In particular, the hidden units in a bidirectional recurrent network are given by the equations

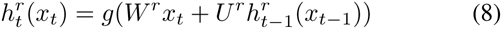

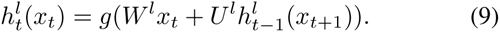

**Fig. 3.**
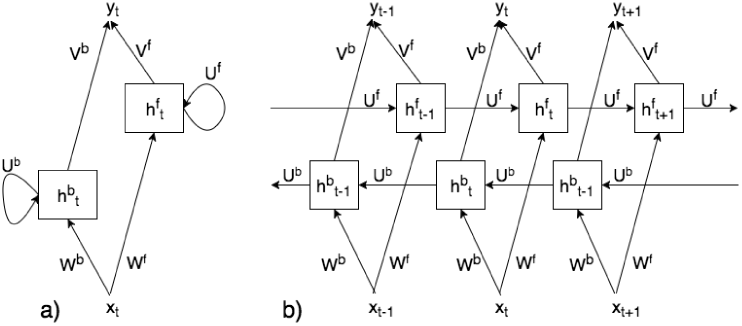
(a) The structure of a bidirectional recurrent hidden unit. (b) A bidirectional recurrent unit unrolled over the time.

We extend the intuition of Siwei et al. [34]. Using a bidirectional recurrent LSTM network we hope to capture the context of the state of a protein within it’s time series. This will allow the machine to have access to information about the past and future of the protein.

### C. Optimization, Parameter Initialization and Regularization

The many non-linear layers in a neural network allow it to become extremely expressive. Unfortunately this large capacity often causes neural networks to overfit before they can learn meaningful relationships. Therefore it is paramount that the model be parameterized to avoid this, especially when trained on a small dataset. Dropout [43] is a simple, yet extremely effective method to prevent feed-forward neural networks from overfitting.

Following the work in [44] which claims that RNNs with dropout do not perform well due to the recurrence amplifying noise, Zaremba et al. [45] propose an implementation of dropout specifically designed for LSTMs. Their proposed method of dropout acts inside the recurrent unit, affecting only non-recurrent connections. This is done by applying dropout only to the values of *x*_*t*_ by replacing *x*_*t*_ by *D*(*x*_*t*_,*p*) in equations (2) - (7), where *D*(*x*_*t*_,*p*) discards node *x*_*t*_ with probability 1 – *p* during each round of training.

Multiple learning optimization methods have been proposed which are applicable to recurrent neural networks. In this word we will use the Adam [46] optimization method computes an adaptive learning rate at each step based on the first and second moments of the gradient. The quality of the local solution is determined not only by the optimization method employed but by the weight initialization as well. The quality of the local solution is determined not only by the optimization method employed but by the weight initialization as well. Glorot [47] introduced Xavier weight initialization as a method to prevent the signals passing through the nodes in a neural network from becoming negligible or unwieldy.

## III. Methodology

### A. Protein Dynamics Datasets

We utilized two largely non-overlapping datasets of proteins from large-scale studies of mutational effects on protein stability. The training dataset of Berliner et al. [21] contains 136 protein structures which were annotated with ΔΔ*G*_*exp*_ of mutation data. The wildtype structures or homologous structural templates used in this dataset were high-resolution. Of these 136 proteins, we excluded all proteins with large chemical cofactors (heme and iron-sulfur clusters) and removed all other chemical cofactors from remaining proteins. A total of 116 proteins were utilized from this dataset. We utilized an additional dataset created by Sahni et al. [9] which contains 950 proteins with both either disease-causing single-point mutations or stable controls. Of this dataset, 884 proteins were utilized after excluding proteins with large chemical cofactors.

Homology modelled was performed for a subset of structures in both datasets where a suitable template was found, as described by Berliner et al. [21]. Molecular models suitable for simulation were constructed automatically using PDB-Fixer [48]. Variable size rectangular simulation cells were constructed for each protein such that there was 8.5 Å of padding with 150 mM of NaCl. All titratable side chains were set to the standard protonatation state at pH 7. Proteins were modelled with the AMBER99SB-ILDN [49] forcefield and water was modelled with TIP3P [50]. All energy minimzation,equilibration, and production simulations were performed with OpenMM 6.31 [51]. All hydrogen bonds were kept rigid and a 2 fs timestep was utilized during equilibration. For production simulations, all bonds were kept rigid and a 5 fs timestep was used. Hydrogen mass was set to 4 amu to facilitate production simulations. For all simulations we utilized a reaction-field electrostatics with a 1 nm cutoff in a periodic simulation cell. Simulations were performed in the NPT ensemble (300 K, 1 atm), with temperature held constant by a Langevin integrator with 1 *ps*^-1^ friction. A Monte Carlo barostat was utilized with a frequency of 25 steps. Data was saved at a frequency of 50ps but all time series were extracted at an interval of 1 ns. The aggregate total simulation data collected for the Berliner and Sahni datasets is 156 *µ*s and 199 *µ*s respectively.

Both datasets had imbalanced proportions of class labels weighted towards destabilizing mutations. Table 1 shows the distribution of stable/unstable class labels for each mutation in the two datasets, and Figure 5 visually represents the distribution of ΔΔ*G*_*exp*_ for the Berliner dataset. As discussed below, we corrected for the imbalanced class labels during machine learning but we did not correct for some proteins being overrepresented in the Berliner dataset.

**Fig. 5.**
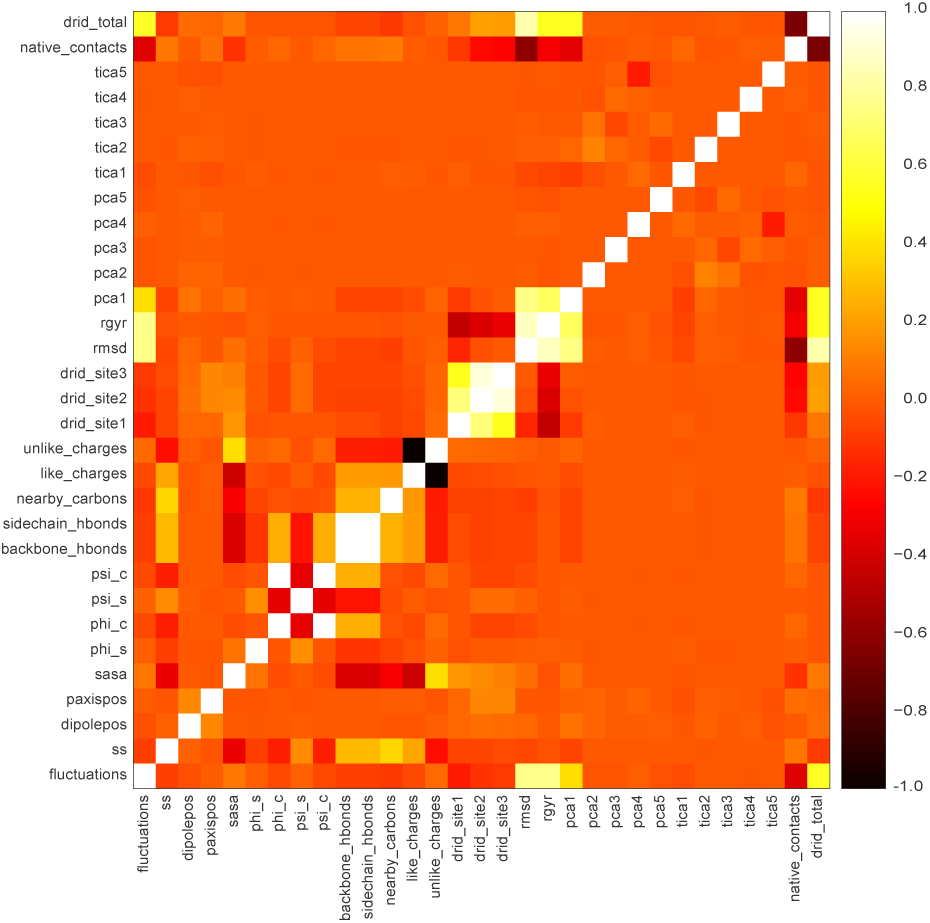
Pairwise Pearson correlation between timeseries features in the Berliner dataset.

**Table I.** DATASET STATISTICS

| Dataset | Stable Mutations | Unstable Mutations | Total Mutations |
| --- | --- | --- | --- |
| Berliner et al. | 710 | 2,208 | 2,918 |
| Sahni et al. | 191 | 1,103 | 1,294 |

### B. Feature Design

Features derived from molecular dynamics simulations were designed to describe the local environment as well as the overall topology of the protein, both of which are poorly described by sequence-based features alone. Berliner et al. computed structural features that quantified the amino acid side chain occupied volume, electrical charge, water accessibility, crowdedness, and amino acid secondary structure. [21] We aim to extract similar features, broadly classified into four types; global timeseries features, mutation timeseries features, static mutation features, and mutation sequence features summarized in Table I, for which each are identified as timeseries features or not. All structural features were extracted using MDTraj [52]

Global features are designed to describe structural properties of the protein as a whole, and include standard stability metrics such as root mean square deviation of all alpha carbons from the initial model and the radius of gyration. The distribution of reciprocal interatomic distances (drid) deviation feature is a similar measure of structural similarity to the initial model, but it provides a better measure of kinetic similarity between structures. [53] These features are complimented by more advanced dimensionality reduction algorithms such as “principal component analysis” (pca) and “time-lagged independent component analysis” (tica) that describe collective motions in the protein. [54] For principal component analysis, C*↵*-C*↵* distances of all residue pairs were utilized, and for time-lagged independent component analysis, all dihedral angles were utilized. Although these global features do not describe the site of mutation alone, they provide supporting information that may help to qualify the degree to which a mutation alters stability as well as flagging important conformational changes that may be occurring in our timeseries.

All mutation timeseries features were extracted at the site of mutation and as such, describe the local environment of a mutation as it would exist in the wildtype form. Traditional residue-specific analysis such as root mean square fluctuations allow for the quantification of site flexibility, something largely absent in static molecular models. Although simulations were conducted with explicit water molecules, they were removed for analysis. As such, we computed a timeries of solvent accessibility using the Shrake and Rupley algorithm as implemented in MDTraj. [52], [55] A timeseries of secondary structure type at the site of the mutation was computed using the DSSP algorithm. [56] Two geometric features were computed to quantify the position of the mutation site alpha carbon with respect to internally defined metrics, the principal inertial axis and the dipole axis. For the computation of hydrogen bonds at the site of the mutation, the Wernet-Nilsson algorithm was utilized. [57] The electrostatic environment was studied by determining the atomic charges within 6 Å of the mutation with respect to the charge of the new side chain being introduced at the mutation site. The backbone phi and psi torsional angles were extracted at the site of the mutation and transformed using sine and cosine to treat the discontinuity at the periodic boundary. The first, second, and third moments of the distribution of reciprocal interatomic distances (drid) was again computed, but this time at the site of the mutation and left in units of reciprocal Å to quantify the crowdedness of the mutation site. [53]

Since simulations were not performed after mutations were introduced, we can not directly estimate how the environment would change upon mutation. However, we can utilize physicochemical data on the change of amino acids at a particular site (”residue change in charge, hydrophobicity, volume, molecular weight”) as a means to approximate this. To compliment these differences in physicochemical empirical data, we utilize a qualitative residue swap similarity metric that is 0 when both the unmutated and mutated amino acids belong to the same class (small nonpolar, small polar, negative charge, large nonpolar, bad behaved, positive charge, side chain amide) and 1 otherwise, as defined by Poultney et al. [58]. Additionally, a static structural feature of potentially high descriptive value is “residue mean mutual information” which is the average value in bits at a particular residue in a mutual information matrix computed using MDEntropy. [59] Finally, two sequence based algorithms are used; the score assigned to the mutation based on the BLOSUM substitution matrix and the score returned by the Provean algorithm for this mutation. [60], [61] In the absence of significant changes of global and site-specific time series, our machine learning algorithms may rely more strongly on physicochemical features to predict stability.

To assist in the interpretation of features, we computed the pairwise correlation between all timeseries in Figure 5. Several groups of features are found to be highly correlated that are intuitively related (residue drid moments, residue backbone and sidechain hydrogen bonds, residue like and unlike charges). Unexpected correlations between residue pairs is also revealed, such as the “first principal component projection” and “root mean square deviation”, as well as the “number of native contacts” and the “distribution of reciprocal interatomic distances”. This analysis suggests that a reduced subset of features might be utilized with minimal loss in accuracy in future studies.

### C. Experimental Setting

The curated protein-sequence datasets are imbalanced, favouring the unstable class. To minimize any bias which this introduces, unstable examples are removed at random from the set of unstable examples until a 45/55 split remains. To mitigate remaining bias, stratified *k*-fold cross validation is used [62]. Specifically we use *nested* 10-fold cross validation. First the data is split into 10 folds and one is selected for the test set. The remaining data is then split into 10 folds and one is selected for the validation set; the rest are used for the training set. Within the inner cross-validation loop, the optimal hyperparameter vector is chosen from candidate set, whose selection is described below. In the outer cross-validation loop the performance of the best-performing models from the inner-folds are assessed on their corresponding test set. The 10 resulting models from this procedure are an estimate of this model’s performance on the entire dataset under the assumption that the 10 models are equivalent to each other allowing us to average their final classification results.

On each of the inner folds the model is evaluated on a candidate set of hyperparameters for validation. Bergstra et al.[63] gave empirical and theoretical evidence that evaluating on randomly chosen hyperparameter vectors is both more efficient and produce better results than most widely used methods of manual search and grid search for initializing parameters. Following this, we empirically choose a set of intervals in which to sample our hyperparameters. The candidate hyperparameters are chosen at random from these intervals during validation.

In the case of an imbalanced dataset the accuracy metric tends to undervalue how well a classifier is performing on the smaller classes. In this case the *F*_1_ score may be a more appropriate metric by which to judge our model. Forman et al.[62] show that several methods of combining *F*_1_-score across folds, including averaging the *F*_1_ score, introduce a significant amount of bias. In the unbalanced class case, missing a single true positive might reduce the *F*_1_ score of a fold significantly, while correctly predicting another true positive has a much less significant impact. Of the methods they test, they found that

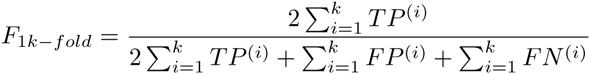

is almost perfectly unbiased. Here *TP*^(*i*)^,*FP*^(*i*)^,*FN*^(*i*)^ are the true positive, false positive and false negative rates for fold *i*. Therefore, this will be the method we use for aggregating *F*_1_- score across folds.

## IV. Classifiers

In this section we outline the methods which we will employ for protein stability prediction.

### A. Bidirectional Recurrent LSTMs

For this problem we have designed a bidirectional recurrent LSTM network. This network can be seen in figure IV-A and unwound across time in figure IV-A. The model consists of two layers of LSTM units followed by a sigmoid activation as output. The first layer of LSTM units is bidirectional, and produces an output at each time step in the many-to-many fashion. The second LSTM layer is only forward-directional and produces a single output at time *T* once all of the timeseries has been read. Dropout and LSTM dropout are applied at various layers throughout the model, indicated by the dotted lines in figure IV-A.

**Fig. 4.**
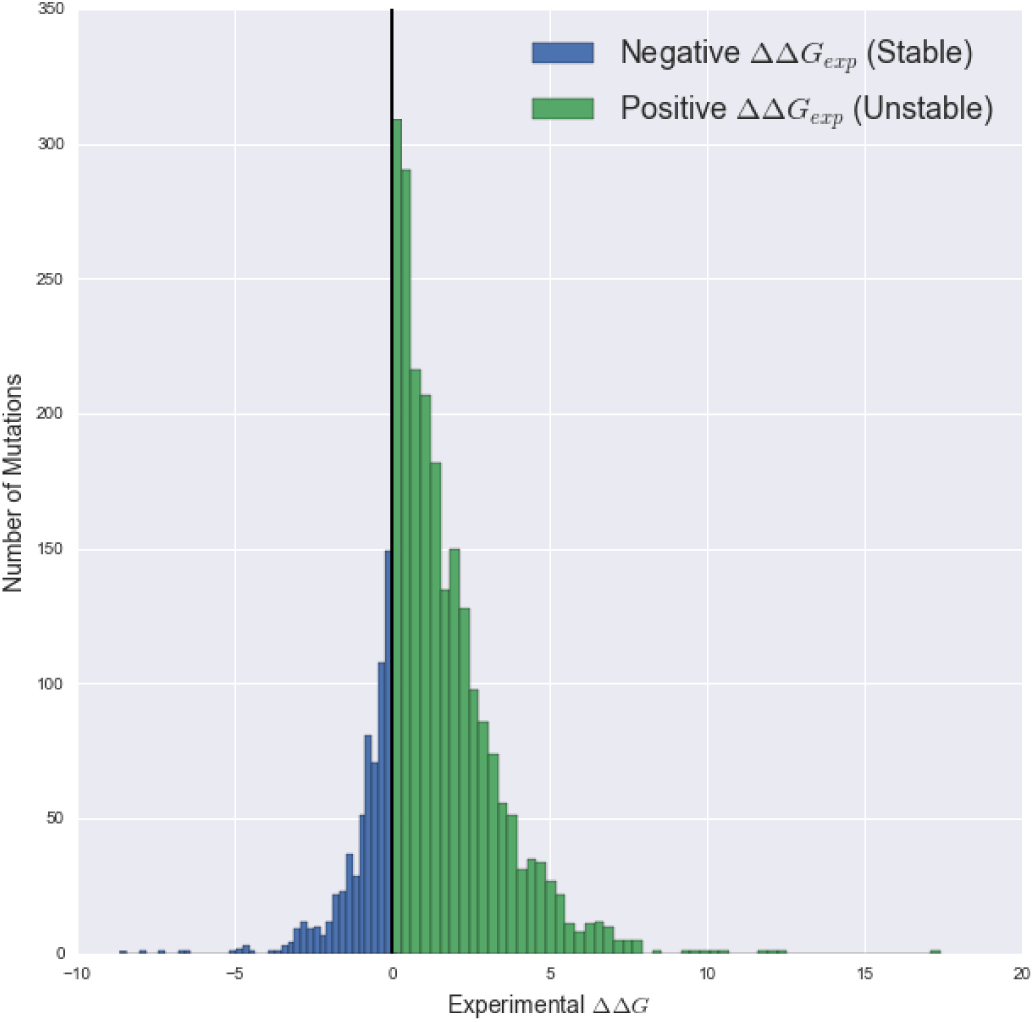
Distribution of ΔΔ*G*_*exp*_ for Berliner et al. dataset

The intuition of the model is that the first layer of bidirectional LSTM units encodes each input within it’s context in the timeseries, while the second LSTM layer reads through these contextualized inputs in chronological order. This can be seen explicitly in figure IV-A.

**Fig. 7.**
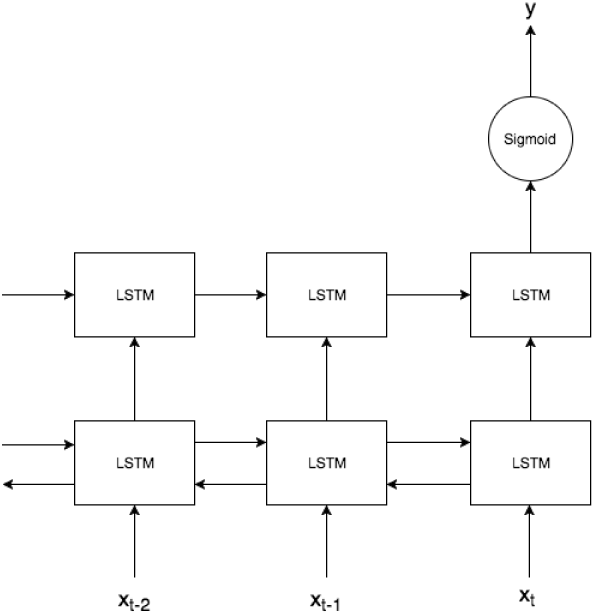
Tuned Bidirectional LSTM Network unwound over time.

The methodology presented in section II-C is used to finetune the model. The weights of each layer are initialized with Xavier initialization, and the bias of the forget gate of each LSTM unit is initialized to 1.0 as suggested in [64]. For training, the Adam [46] method is used along with binary cross-entropy as the loss function. We observed empirically that the model performed optimally when the first bidirectional LSTM connection contained 11 nodes each for the forwards and backwards connections, the second LSTM layer contained 5 nodes, and the sigmoid layer contained only a single node.

During cross-validation our model tended to perform better when dropout to the layer of bidirectional LSTM nodes was fairly high, *p* ≈ 0.6, dropout to the regular LSTM layer was lower, around *p* ≈ 0.45, and dropout before the sigmoid output was quite small at *p* ≈ 2. Better results were seen with smaller step size *α* ≈ 0.001, and large values of Adam decay parameters *β*_1_, *β*_2_ õ [0.9, 1).

For comparison we also evaluate two simpler recurrent neural networks: A standard recurrent neural network with a single layer of 15 recurrent nodes followed by a sigmoid output, which we will call the *RNN model*. And a simpler form of bidirectional LSTM with only a single layer of bidirectional LSTM nodes in the many-to-one fashion, with 15 LSTM units per direction. This is followed by a single sigmoid output. We call this model the *simple Bidirectional LSTM model*. Both of these networks employ dropout between layers, have weights initialized with Xavier initialization, and are optimized with Adam with a binary cross-entropy loss function, in the same fashion as was done for our tuned bidirectional LSTM model above. The neural networks were implemented using the Theano [65] and Keras [66] packages.

### B. Simple Machine Learning Models

Seven simple machine learning models were used to benchmark the relative success of our neural networks. Models were tuned using an iterative grid search to find the optimal hyperparameters. We utilized the Gaussian Naive Bayes model, k-nearest neighbours, support vector machines, stochastic gradient boosting of decision trees, random forests, and AdaBoost. Note that the objective of this work was not to compare the strengths and weaknesses of each of these approaches on our dataset.

## V. Results

### A. Protein Dynamics Example

There are 553 single-point mutations to the micrococcal nuclease protein (PDB: 4WOR) in the Berliner dataset. This is a bacterial protein is an enzyme that breaks apart single-stranded nucleic acids. A structure of this protein has been available since 1969, making it an extremely well-studied protein with a great deal of experimental data available regarding its thermostability upon mutation. We performed 1.6 *µ*s of simulation for this protein (in the absence of any nucleic acids) and examined any structural fluctuations of the entire protein, as well as site-specific information related to regions with ΔΔ*G* of mutation information. Here we present several timeseries related to a specific mutation G83W in Figure 8. Since Gly is an unusual amino acid without a side chain and Trp is a large nonpolar residue, one might expect that this is a destabilizing mutation, and indeed this is found to be destabilizing in the ProTherm database. However, the Provean algorithm predicts this mutation to be be neutral. We expect that the timeseries extracted in at residue 83, such as the the root-mean square fluctuations and change in number of like charges will assist in correctly classifying this mutation as destabilizing. Interestingly, one may notice that transitions are not made between multiple basins in the space deposition timeseries of time-lagged independent component 1 and even principal component 1, suggesting that it is likely that we have not obtained sufficient sampling of this protein along both its slowest and highest-variance degrees of freedom. Nonetheless, we obtain additional info

**Fig. 8.**
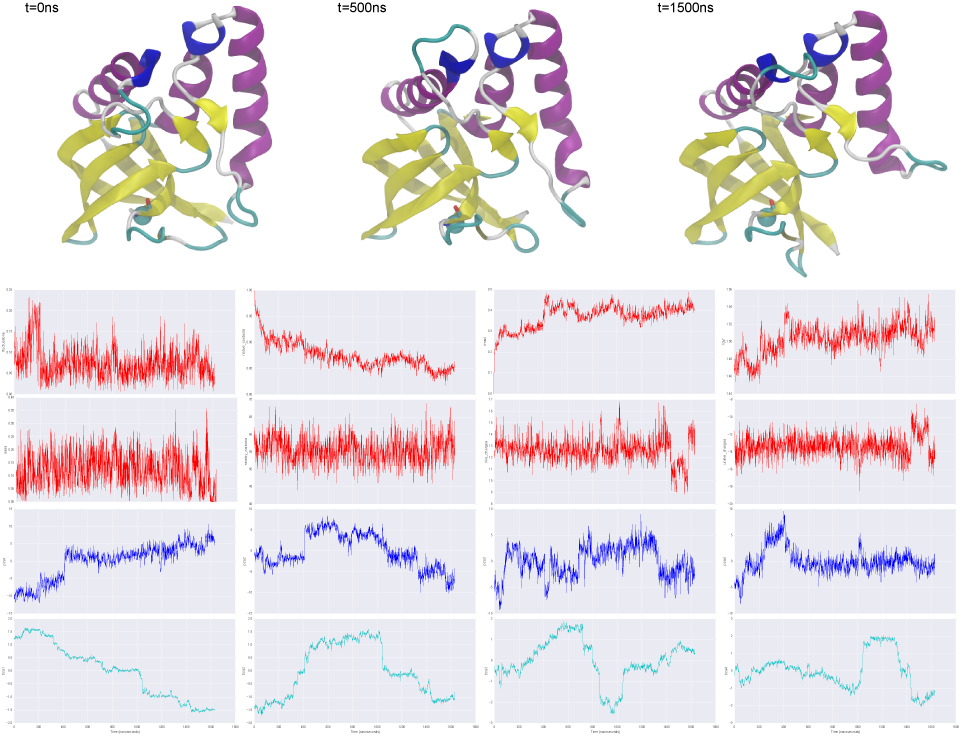
Micrococcal nuclease protein molecular dynamics example. (top) Molecular renderings of this protein throughout the timeseries colored by secondary structure. (bottom) Selected feature timeseries related to the mutation G83W where the alpha carbon of residue 83 is shown with a cyan sphere. From top left to bottom right; fluctuations, native contacts, rmsd, radius of gyration, solvent accessible surface area, nearby carbons, nearby like charges, nearby unlike charges, principal component 1-5, time-lagged independent components 1-5.

### B. Machine Learning Benchmarks

We report the accuracy and F1-score of the six basic machine learning models studied using mean and variance features of our timeseries data in Figure 9. We achieved the worst performance using the gaussian naive Bayes classifier for both scoring metrics on both the Berliner and Sahni datasets. The most successful of models were ensemble methods, gradient boosting, random forest, and AdaBoost classifiers, all of which were nearly within error bars and ranged between 64% and 75% accuracy for the Berliner dataset and between 70% and 76% accuracy for the Sahni dataset. In general, F1-scores were consistently lower than our accuracy scores, and we expect they represent a fairer representation of the performance of our models. We observed marginally lower performance using histogram binned features in the Berliner dataset, ranging from 0% to 10% across all models. This suggests that both dimensionality reduction techniques we applied to our timeseries features resulted in effectively the same results. Our highest performing models across both datasets were the “gradient boosting” and “random forest” ensemble methods using the mean/variance features. Using mean/variance features, these two models achieved an accuracy of 71% ± 4% and 75% ± 4% respectively on the Berliner dataset. The same models achieved an accuracy of 76% ± 3% and 76% ± 2% respectively on the Sahni dataset.

**Fig. 9.**
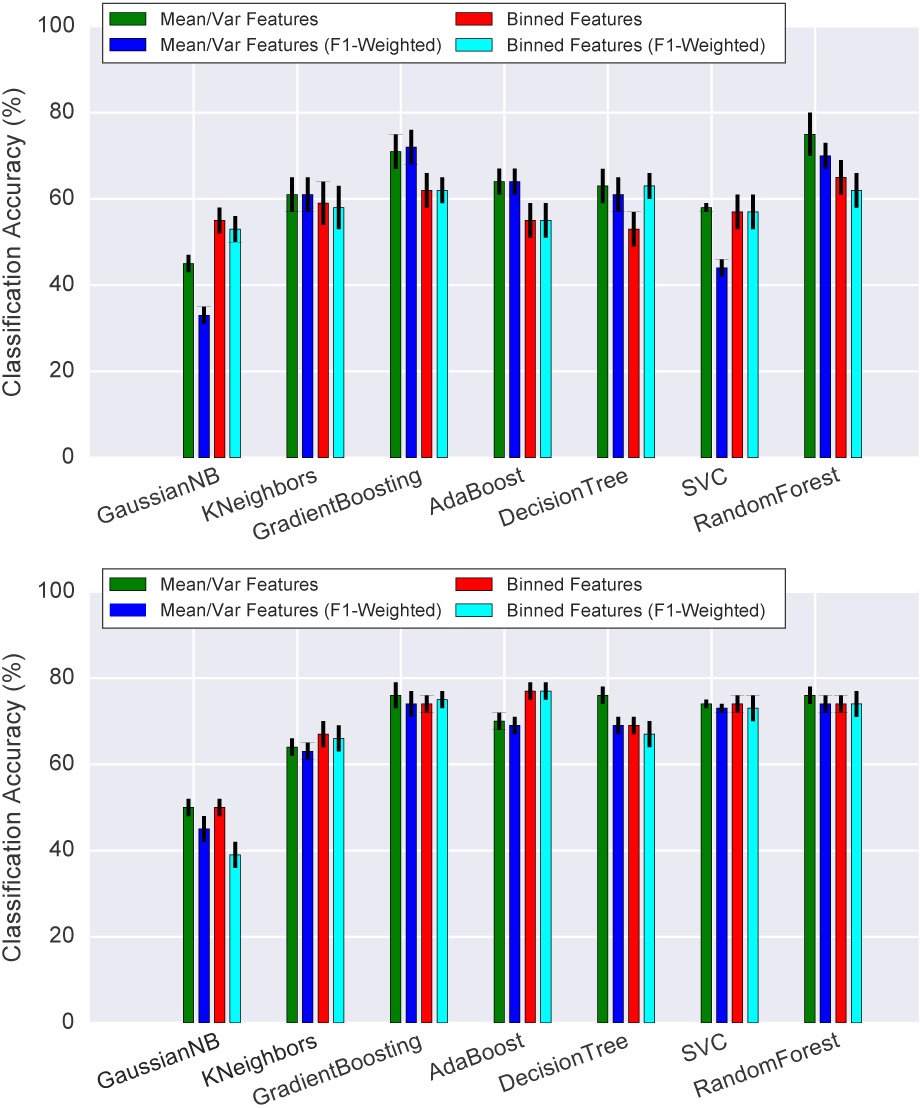
Accuracy of supervised machine learning algorithms using mean/var and histogram binned timeseries features (top). Classification accuracy and F1-scores for the Berliner dataset (bottom). Classification accuracy and F1-scores for the Sahni dataset. Error bars are computed using the standard error of mean over 10 folds.

### C. Recurrent Neural Network

We report the accuracy and F1-score of the three recurrent neural network models studied using timeseries data in Figure 10. It was found that both a bidirectional LSTM offered the highest accuracy and F1-score for the Berliner dataset, 76% and 80% respectively. The same bidirectional RNN obtained an accuracy and F1-score of 73% and 75% when trained on the Sahni dataset. The simple bidirectional LSTM and recurrent neural network obtained only marginally lower scoring metrics. Here we experimented with the removal of sequence based features during the training of the neural network. Our results suggest that sequence features resulted in a gain of classification accuracy of 2% to 3% but do not appear to be required for classification. Similarly, we examined the accuracy of ternary classification (destabilizing, neutral, and stabilizing groups), although due to a limited number of class members, the performance of this method dropped by approximately 25% for all recurrent neural networks.

**Fig. 10.**
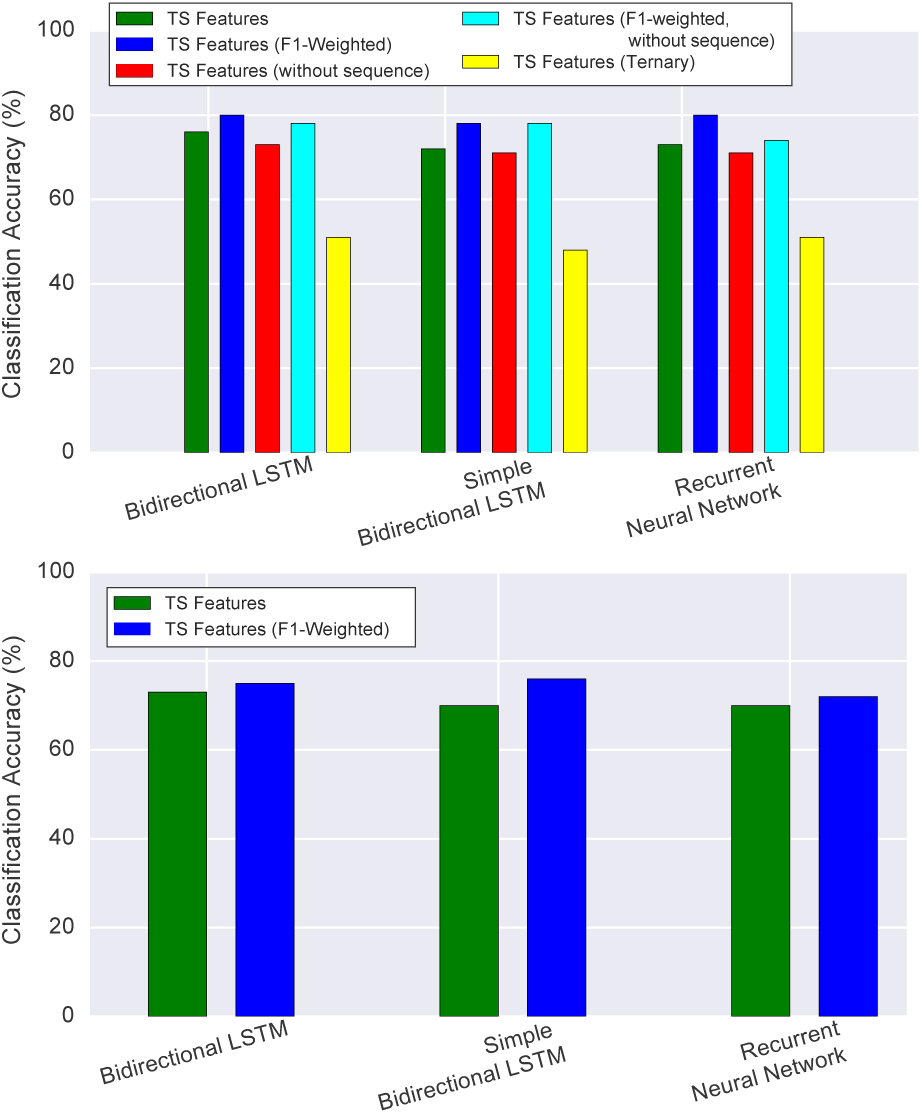
Accuracy of several recurrent neural network types utilizing complete timeseries features (top). Classification accuracy and F1-scores, with and without sequence features for the Berliner dataset (bottom). Classification accuracy and F1-scores for the Sahni dataset.

### D. Comparison to Existing Methods

In order to assess the quality of our models, we compared to several high-performing stability prediction algorithms in the literature as summarized in in Table II (ELASPIC [21],our dataset. The three former methods were designed to not only to classify mutations based on structure, but to predict ΔΔ*G*. As such, we imposed ΔΔ*G* cutoffs consistent with our methodology to draw comparisons between our class predictions and these regression and energy-function based methods (< 0 for neutral, ≥ 0 for deleterious). Berliner et al. authored the ELASPIC methodology and subsequently used the Berliner dataset for training, we note that it had the highest performance on this dataset using accuracy and F1-score as metrics (73% and 80%). The VIPUR methodology was found to be high performing on the same dataset. The ΔΔ*G* predictions of the FoldX algorithm, which was also utilized within ELASPIC and found to be among the highest performing features, had only slightly worse accuracy the ELASPIC algorithm. Similarly, the Provean score (which was classified as neutral or deleterious based on a cutoff of - 2.5) was also used in the ELASPIC algorithm as a feature, but it is frequently used by itself to assist in the prediction of mutation stability by itself and thought to have higher accuracy than popular PolyPhen2 algorithm. [61] It is not unexpected that all algorithms perform poorly on the Sahni dataset since they were trained using ΔΔ*G* values and the stability metric used by Sahni et al. is considerably different. Note that we did not attempt to train our models on the Berliner dataset and classify stability of the Sahni dataset. As such, we would expect a comparable drop in performance. The VIPUR algorithm was not run on the Sahni dataset and will be better assessed in future studies. To summarize, our best classification algorithm appears to be equal or superior to the majority of these approaches.

**Table II.**
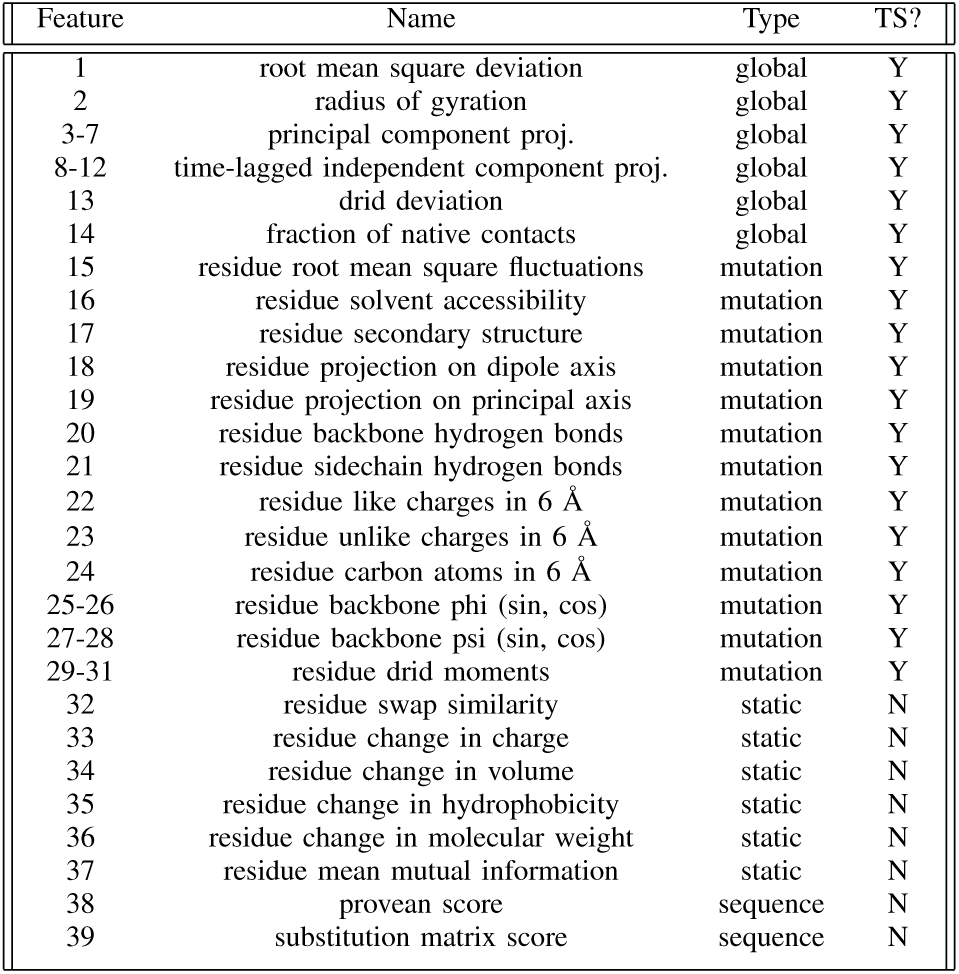
MACHINE LEARNING FEATURES

**Table III.** ACCURACY COMPARISON TO EXISTING METHODS

| Methodology | Berliner Acc. | Sahni Acc. | Berliner F1 | Sahni F1 |
| --- | --- | --- | --- | --- |
| ELASPIC | 73% | 42% | 80% | 37% |
| VIPUR | 45% | - | 63% | - |
| FoldX | 71% | 40% | 80% | 38% |
| Provean | 64% | 35% | 75% | 39% |

## VI. Visualization and Feature Importance

### A. Garson’s Method For Recurrent Neural Networks

Several methods have been proposed to determine the importance of the input nodes to the neural network [67], [68]. Unfortunately none of the presented methods are immediately suitable for handling recurrent connections, and few of them have been generalized beyond a single-layer neural networks. A simple method for evaluating the importance of the inputs was proposed by Garson [69], extended by Goh [70], and Gevrey et al. [68]. Garson’s algorithm phrases the importance of an input as the sum of the weight of the (directed) paths through the neural network from that input to that target. Let *N* be a neural network with *n* input nodes, a single layer of *m* hidden nodes, and *k* target nodes. Garson’s algorithm states that the importance of input node *x*_*α*_ is

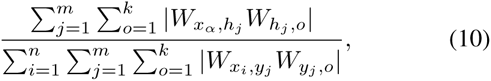

where *W*_*ij*_ is the weight between node *i* and node *j* in the neural network. If there is no edge between nodes *i* and *j* then *W*_*ij*_ = 0.

We show how this can be generalized for recurrent units. To extend this to arbitrary-depth neural networks, we can rewrite equation 10 by defining the *relative importance* of a node *x*_*α*_ in the neural network to be

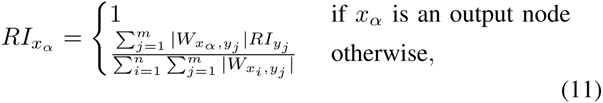

where *n* is the number of nodes on the same layer as *x*_*α*_ and *m* is the number of nodes on the layer which *x*_*α*_ has outgoing edges to. The relative importance of a node is equivalent to summing over the product of the weights along each path between *x*_*α*_ and any of the output nodes.

It is straightforward to extend this recursive definition to the simple recurrent units in equation 1, figure II. This is done by unwinding the recurrence. Consider the simple structure in figure II with a single layer of recurrent units *h*_0,1_,…,*h*_0,*n*_,a layer of inputs *x*_1_,…,*x*_*m*_, a layer of outputs *y*_1_,…,*y*_*k*_,and weights *W*_1,1_,…,*W*_*m,n*_ between *x*_*t*_ and *h*_*t*_, *V*_1,1_,…,*n,k* between the hidden units *h* and the output *y*s, and recurrent weights *U*_1_,…*U*_*n*_. The relative importance of the recurrent unit *h*_0,*j*_ can then be found by expanding equation 1,

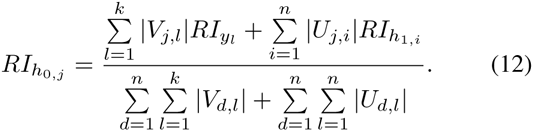

Note that here we are taking the relative importance of each input node at time *t* = 0, as by the equation 11, this allows us to capture the full time dependencies across *t* = 0,…,*T*.

This method can in theory be extended to LSTM units, but calculating the recurrent relation becomes far more involved. Therefore we evaluate the the importance for each of the features for our RNN model, and leave the calculation of the relative importance of LSTM units for further work.

It is important to note that unlike equation 11 for standard feed-forward neural networks, summing over each *RI*_*h*_*0,j*__ for *j* = 1,…,*n* does not sum perfectly to 1 for finite *T*, although for *T* → ∞, summing over each *RI*_*h*_*0,j*__ for *j* = 1*,…, n* does converge to 1. The impact of each hidden unit *h*_*t,j*_ on *RI*_*h*_*0,j*__ as *t* grows decreases exponentially in the weights *U*_*ij*_. Therefore this relation can be accurately approximated by taking large enough *T*.

We calculate the relative importance of the input features to the RNN of the for *T* = 20. This value of *T* was chosen because it becomes computationally intractable for much higher values of *T*, and the changes in the relative importance are negligible. The results can be seen in Figure VI-C, and are discussed in section VI-C.

### B. Neural Interpretation Diagrams

We provide a modification of the model of neural interpretation diagram as presented in [71]. Neural interpretation diagrams (NIDs) provide a way for us to visualize the effect one node has on another in a neural network. In a NID each row of circles correspond to a node in a layer in the neural network. The nodes are connected by edges, representing the weight between those two nodes. An edge is grey if the associated weight is negative, and black if positive. The width of the edge reflects the magnitude of that weight.

To remedy this, we propose a modification of neural interpretation diagrams. Two contrasting colours are chosen to coloured cyan, while negative weight edges are magenta.represent the edges. Edges which have positive weight are than those with low weight. To handle the issue of NIDs causing nodes to appear more important than they truly are, we combine our extension of Garson’s method to NIDs. The size of each node in the NID is now dependent on the relative importance (equation 11) of that node within the network. As well, we colour each node is the normalized sum of the incoming weights to that node. The colour of the input nodes is the normalized sum of the outgoing weights from that input. As biases do not receive a relative importance, their size is the sum of the magnitude the weights which are connected to it.

Furthermore, we extend NIDs to recurrent neural networks. A separate NID is used to plot the recurrent connections for the network. This can be seen for our recurrent neural network in figure VI-B. Rather than using the the relative importance scores for the size of the nodes in the recurrence NID (figure VI-B (b)), we believed that the sum of sum of the incoming or outgoing weights would be more informative.

**Fig. 6.**
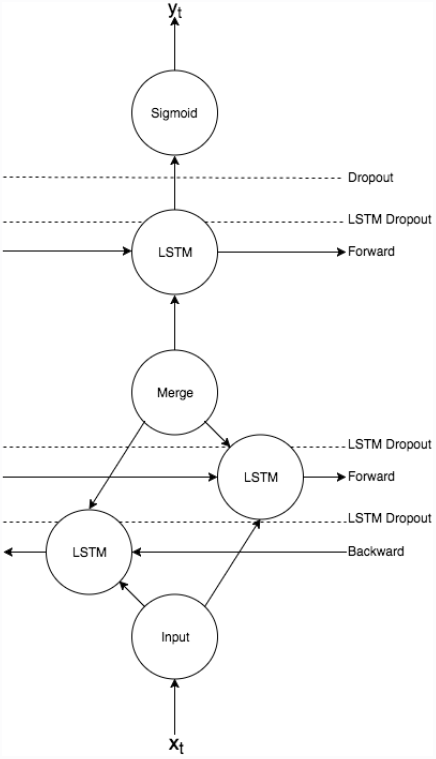
Structure of tuned bidirectional LSTM neural network.

Finally, we outline how to use the above multi-NID method to extend NIDs to LSTM recurrent neural networks. The feed-forward portion of the network is graphed as above. Due to the clutter of LSTM units, we use two plots to display the connections in the LSTM unit. In the first, the feed-forward weights are displayed, while in the second the recurrent weights are displayed. The colour and represent the same features as described above for the RNN. The size of each node now represents the magnitude of the sum of the incoming weights to that node, rather than it’s relative importance because the recurrence for the LSTM unit became unwieldy to calculate. The size of the input nodes represent the magnitude of the sum of the weights leaving that node.

The NID for our tuned bidirectional LSTM model can be seen in figures VIII, VIII VIII,VIII,VIII,VIII,VIII in the supplementary information. Unfortunately we found that our extended NID without the extra information provided by Garson’s method did not simplify the representation of the regular NID in order for us to analyze analyze coherently. Therefore we focus on the RNN for which we can implement Garson’s method for our feature analysis, relying on the fact that they differ only slightly in performance. This exemplifies why including Garson’s method (equation 11) is extremely useful for NIDs.

### C. Feature Discussion

We analyze the NID for the RNN presented in figure VI-B, and the results from Garson’s method which are shown in the first histogram in figure VI-C. If a feature has predominantly positive (cyan) paths leading from it, then a large value for that feature contributes to the neural network predicting the positive class (unstable). On the other hand, if the feature has many negative (magenta) paths leading from it, then a high value of that feature contributes towards it predicting the negative class (stable). We say that such nodes have high negative or high positively weight respectively.

**Fig. 11.**
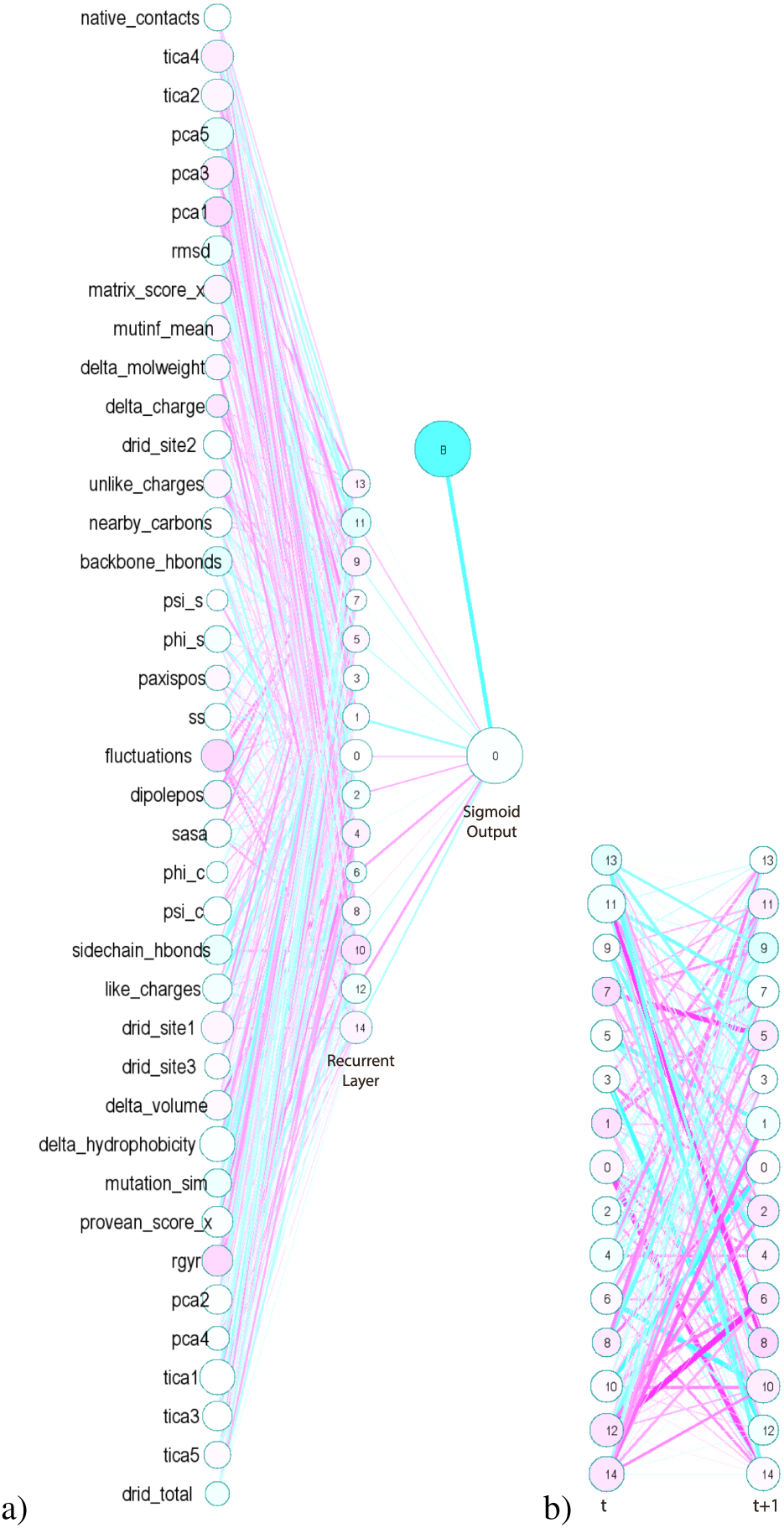
A modified NID for our RNN model. a) The feed-forward connections of the neural network. b) The weights of the recurrent connections of the neural network at time *t* and *t* + 1.

Nodes with high positive weight include “number of sidechain and backbone hydrogen bonds”, as well as “mutation similarity score”. Both of these features are expected to be essential for characterization of the effect of mutation, although interestingly the latter is a static value. Nodes with high negative weights include “principal component 1 projection”, “root mean square fluctuations”, and “radius of gyration”. Interestingly, all of these were global features thought to strongly characterize the stability of the protein. The determination of highly weighted nodes in our RNN provide motivation for the development of even more robust features to assist in classification accuracy. An example of this would be a more advanced measure of hydrogen bonding involving the residue at the site of mutation, further breaking down the electrostatic properties in this environment in a similar way to the energy terms returned by software like Rosetta and MODELLER. This may also draw attention to the limitations of NIDs for visualizing neural networks.

Figure VI-C shows the relative importance (multiplied by 1,000) of features to the neural network as calculated by our extension of Garson’s method. The differences in feature importance was not nearly as significant as was seen for the other ensemble machine learning methods plotted in the lower figures, although similar trends were observed. In particular the change in hydrophobicity appeared as a significantly important feature for many methods. While the mean fluctuations at the site of the mutation is understandably a high importance feature due to the potentially stable or unstable environment of the mutation, but the recurrent neural network also found the “time independent component analysis project 1” and “principal component analysis 1” to be of high importance. The mean and variance of these features have negligible information so it is reassuring that the neural network was able to utilize this dynamic structural information. As these specific features cannot be determined from any existing structural, sequence, and energy function based method, it is reassuring that the neural network was able to utilize them effectively for classification.

Relative feature importance is also presented for several ensemble machine learning algorithms in Figure VI-C. Unlike the uniform neural network feature importance values, several algorithms were found to rely heavily on individual features. The consensus across ensemble machine learning algorithms is to put high importance on static features described physicochemical properties of amino acids involved in the mutation. Since we do not explicitly model the presence of mutated amino acids, we rely strongly on these features to characterize the mutation. Amongst our top performing ensemble methods, random forests and gradient boosting, several mutation site specific features were found to have high importance, including the mean solvent accessible area and the mean and variance of the number of like charges and root mean square fluctuations. The relatively low important of global properties suggest that we may have poorly described the overall topology and stability of the protein with our global features. This analysis of feature importance draws attention to limitations of our featurization and reveals areas of improvement for feature engineering for the prediction of stability.

## VII. Discussion

Both the simple machine learning models used in this manuscript as well as our top performing model (the bidirectional LSTM) were highly effective at the classification of arbitrary protein mutations as neutral or deleterious. Our results support that an optimized bidirectional LSTM network with dynamic timeseries features is capable of surpassing simple machine learning algorithms, but not by a large degree. The complexity of the recurrent neural network, both in implementation and interpretation of the output suggests that its use may require additional work for applications such as this one. Improvements to this approach may include further optimizations of neural network architecture and hyperparameters.

**Fig. 12.**
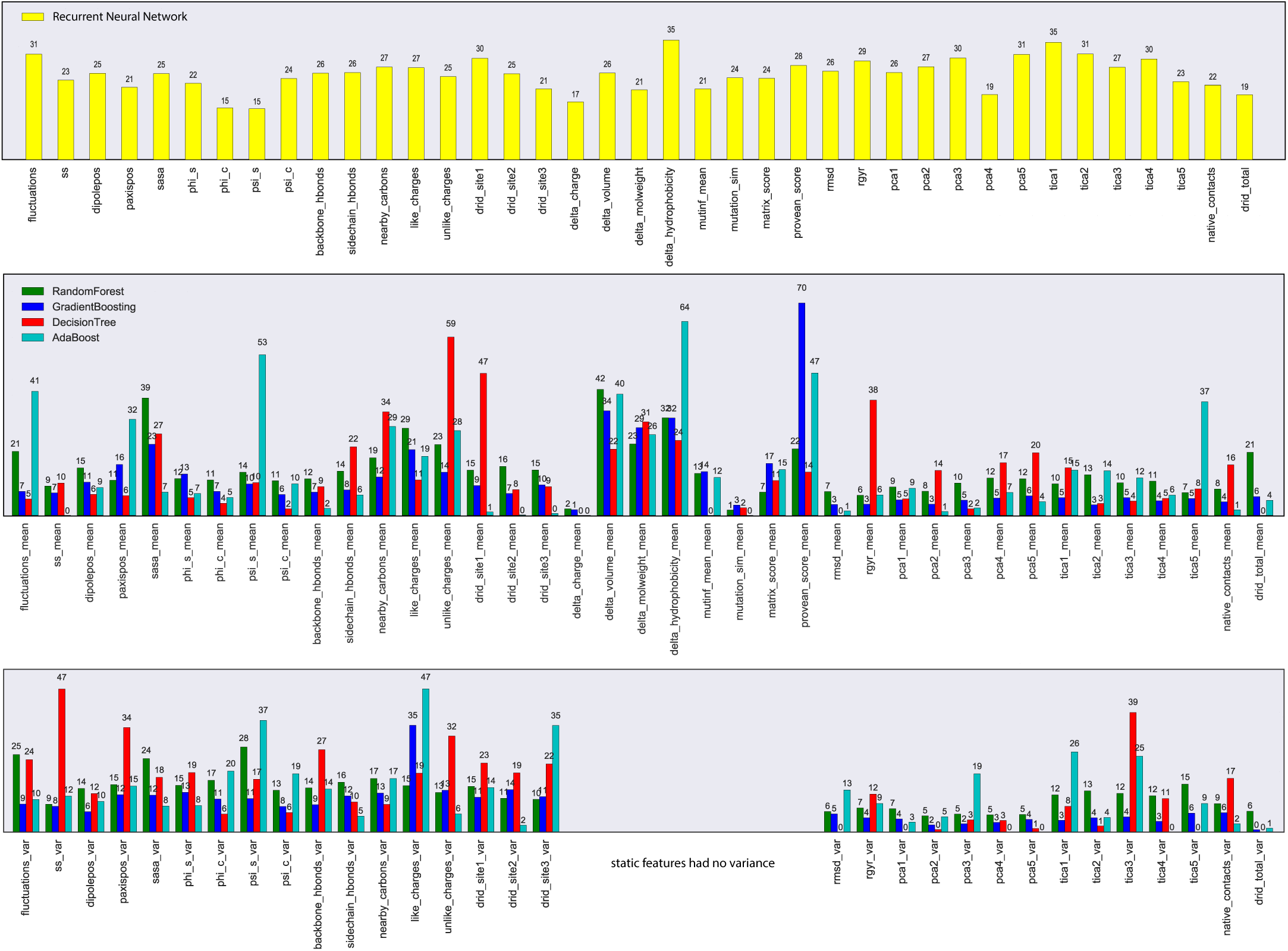
Feature importance histograms for the recurrent neural network and ensemble machine learning models. (top) The relative importance (multiplied by 1,000) of timeseries features in the recurrent neural network model. (middle) The relative importance of mean features used with several ensemble-based machine learning algorithms. (bottom) The relative importance of variance features used with several ensemble-based machine learning algorithms.

Even though this study utilizes one of the largest atomistic multi-protein molecular dynamics datasets, with comparable size of Dynameomics database in terms of aggregate simulation for our two datasets [72], we expect that many more proteins must be simulated to achieve higher performance using a recurrent neural network. We expect that potentially an order of magnitude more proteins (and a similar number of labelled examples), may be required. Using todays computing resources, this may be achieved by accelerated simulation sampling algorithms like simulated tempering at the expense of losing actual dynamics. [73]. Although long simulations (up to 2 microseconds) were computed, it cannot be ignored that long timescale domain reorganization of proteins may still occur, as seen in long time-scale simulations of BPTI [74]. Even so, it is difficult to assess if longer protein simulations would be required in order to improve classification accuracy of our neural network. Additional testing, potentially involving repeated truncation of timeseries data and retraining, would need to be performed in order to assess if our simulations are sufficient in length. Nonetheless, the dynamic dataset generated for this study will likely be a highly valuable resource in the development of hybrid methods that utilize structure, dynamics, sequence, and energy functions.

In future studies we hope to test the methodology presented in this manuscript using other databases of known disease-causing mutations such as OMIM [75], HAPMAP [76], and COSMIC [77]. Testing on these databases of mutations will inherently require simulations of even more proteins to extract timeseries features. As our dataset of simulated proteins increases, it becomes increasingly important to study if it is acceptable to train our model using more than one datasets even though they may not share a common stability metric (ΔΔ*G*, pathogenicity, ternery labelled points). It is also not known if our recurrent neural network approach can be modified to perform regression, and potentially make quantitative ΔΔ*G* predictions. This would greatly increase the significance of our approach and make it easier to compare to other regression-based algorithms.

From a broader perspective, three major factors restrict the applicability of our method to rapid clinical diagnostics. Firstly, the generalization of our approach to a proteome-wide scale cannot be assessed because homologous structures are not known for a large percentage of human proteins. However, as new structures and structure determination methods are discovered this may change. Secondly, a fundamental limitation of our methodology is that it lacks proper treatment of interfaces (sites of protein-protein, protein-DNA, protein-RNA, protein-ligand, and protein-cofactor interactions). As it was determined from Sahni et al. [9], many disease-causing mutations do not significantly alter protein folding and stability, but rather protein-protein interactions. As databases of protein interaction sites and site prediction algorithms become more robust, these factors may be included as timeseries features for this approach, but for now, this represents a significant limitation to the connection of mutation deleteriousness and disease. Finally, our method requires long-timescale simulations to be performed on all wildtype proteins in training and test datasets, potentially requiring months or years of simulation. It is possible that simulations may eventually be precalculated on a large subset of all human proteins in the protein databank, but this likely represents years of continuous computation and will require collaboration of multiple simulation labs.

This work presets novel research regarding the use of dynamic structural features for mutation stability prediction. However, additional experiments and validation are required before a tool such as this this could be used for applications like protein engineering through thermostability optimization or clinical diagnostics.

## ACKNOWLEDGMENT

The authors would like to acknowledge the support of Alexey Strokach and Dr. Philip Kim from whom we obtained both the Berliner and Sahni datasets, along with helpful discussions related to this report. We are grateful for useful discussions and support from Dr. Régis Pomès. We acknowledge the support of CPU computing resources on the Parallel supercomputer provided by WestGrid, GPU computing resources on the Helios supercomputer provided by Calcul Quebec and Compute Canada.

